# BrainSTEM: A multi-resolution fetal brain atlas to assess the fidelity of human midbrain cultures

**DOI:** 10.1101/2024.09.17.613274

**Authors:** Hilary SY Toh, Lisheng Xu, Carissa Chen, Pengyi Yang, Alfred X Sun, John F Ouyang

**Affiliations:** Duke-NUS Medical School, Signature Research Program in Neuroscience and Behavioural Disorders, Singapore, Singapore; Computational Systems Biology Unit, Children’s Medical Research Institute, Faculty of Medicine and Health, University of Sydney, Westmead, NSW 2145, Australia; Charles Perkins Centre, School of Mathematics and Statistics, University of Sydney, Sydney, NSW 2006, Australia; Department of research, National Neuroscience Institute, Singapore, Singapore; Duke-NUS Medical School, Centre for Computational Biology, Singapore, Singapore; Duke-NUS Medical School, Signature Research Program in Cardiovascular and Metabolic Disorders, Singapore, Singapore

**Author notes:** For correspondence (AXS); (JFO). These authors contributed equally to this work (Hilary SY Toh, Lisheng Xu).

## Abstract

Many midbrain dopaminergic neuron (mDA) differentiation protocols aimed at Parkinson’s disease (PD) modeling and cell replacement therapy have been developed. However, comprehensive evaluations of the transcriptomic fidelity of these protocols at the single-cell level against a common *in vivo* reference have been lacking. To this end, we constructed an integrated human fetal whole-brain atlas and a midbrain subatlas to use as a standard of comparison. From the whole-brain atlas, we observed distinct brain-region-specific gene expression in most neural cell types, emphasizing the need to first evaluate *in vitro* protocols at the whole-brain level to identify midbrain-associated cells. These cells are then mapped to the midbrain subatlas for more refined neuronal subtype specification and trajectory analysis specific to the midbrain. We surveyed all publicly available single-cell datasets of human midbrain culture models and performed the two-tier mapping. Using this biologically-driven multi-resolution mapping strategy which we termed BrainSTEM (brain Single-cell Two tiEr Mapping), we confirmed the presence of multiple midbrain cell types (‘on-target’), but also a substantial proportion of cells associated with non-midbrain regions and subtypes (‘off-target’). This leads to an overall ‘inflation’ of mDA presence, stemming from non-midbrain-associated cells, across all published protocols. BrainSTEM thus offers an unbiased framework for understanding the current state of midbrain models and aids the improvement of midbrain differentiation protocols for PD studies.

## Introduction

Single cell transcriptomic technologies have enabled an unprecedented growth in our understanding of the human brain spanning from fetal development to the adult brain, in both healthy and disease states. Neural cultures have also advanced from two-dimensional neuron cultures to 3D organoids and even multi-region combinations of organoids termed assembloids^1^. Many groups utilize single-cell RNA sequencing to assess the fidelity of their neural models by comparing their transcriptomic similarity to the brain. To this end, much attention has been directed towards cortical models^2,3^, especially with the advancement in cerebral organoid models^4,5^ in recent years. To date, many midbrain differentiation protocols^6-10^ have been developed but remain relatively less well-characterized. Furthermore, there have not been comprehensive surveys to characterize the cell states found in midbrain models and to systematically compare available midbrain datasets to the *in vivo* midbrain.

The human midbrain has been the target of numerous studies due to the selective vulnerability of midbrain dopaminergic (DA) neurons in Parkinson’s disease (PD). Many DA neuron differentiation protocols have been proposed, modeled after developmental biology principles to pattern stem cell derived progenitors towards a ventral floor plate fate using small molecule morphogens such as sonic hedgehog (SHH) and WNT agonists. While midbrain differentiation protocol development continues to undergo refinement, considerable variation in culture conditions persists. Thus, it remains a challenge to assess how well each protocol recapitulates the human midbrain, and by extension how effectively it can serve as a model for PD or a basis for cell therapy products for transplantation. In a disease modeling context, the variability in midbrain protocols may complicate the interpretation of disease phenotypes and hinder the development of therapeutics and biomarker identification. The heterogeneity in DA neuron yield and uncertain transcriptomic fidelity of culture products also poses challenges for PD cell therapy, an active area of clinical trial research^11-13^.

Commonly used methods to assess midbrain differentiation models involve immunostaining or quantitative real-time PCR (qRT-PCR) of single gene markers such as tyrosine hydroxylase (TH) to label target cell types like DA neurons. This approach is limiting as each marker gene may mark several neuronal populations, resulting in an overestimation of the target cell type yield. Using single cell approaches offers a more global view of the transcriptomic landscape of midbrain models. However, a frequent pitfall is comparing midbrain protocols primarily to fetal midbrain references instead of the whole brain, which prematurely limits the scope of the analysis and can introduce bias as this comparison may not account for the presence of cell types from other brain regions found in midbrain models due to off-target patterning.

In this study we address these questions by developing a comprehensive and multi-resolution fetal whole brain atlas with a focus on the midbrain and characterize the physiological fidelity of an extensive selection of available midbrain culture datasets. The fetal midbrain atlas is designed to provide a detailed understanding of the developmental trajectory of mDA neurons, helping to elucidate the neuronal progenitors that give rise to DA neurons. To explore the variability among midbrain differentiation protocols, we conducted an extensive curation of available single-cell transcriptomic datasets of the midbrain derived from a variety of protocols that cover both two-dimensional and 3D organoid culture models. Recognizing that *in vitro* differentiation inevitably produces additional cell types and regions other than those desired, we designed a two-tier projection approach that retains midbrain-associated cell types to evaluate the fidelity of midbrain protocols to the human midbrain at single cell resolution. In the first tier, query datasets are projected onto the fetal whole brain atlas to assess the brain region specificity of the various neural populations. Following this, midbrain-specific cells are further projected onto a more intricately annotated midbrain subatlas. This second, more nuanced, projection allows us to uncover midbrain DA neurons as well as other rarer ‘off-target’ subpopulations that would otherwise be masked by the presence of non-midbrain region cell types. Using this two-tier mapping approach which we termed BrainSTEM (Brain Single-cell Two tiEr Mapping), we examined each protocol’s capacity to recapitulate the *in vivo* midbrain in terms of their cell type diversity, proportion and transcriptomic similarity. Collectively, our work offers comprehensive whole brain and midbrain atlas resources and introduces a novel framework for evaluating the accuracy of midbrain models to optimize protocol development going forward.

## Results

### Integrated whole brain fetal atlas identifies brain region-specific cell types and associated gene signatures

To create a fetal whole brain reference atlas containing brain region information, we integrated two datasets published recently by Braun *et al*.^14^ and Zeng *et al*.^15^ (Figure 1A). We leverage on the advantages of both datasets to profile the human brain from post-conception week (PCW) 3 to 14 totalling 679,666 cells from 39 donors, providing a highly comprehensive view of early human brain development. The total number of cells is less than that from both studies combined as we sampled the same number of cells for each donor before carrying out batch effect correction according to the reference papers’ methodology (Methods). Doing so enables us to preserve biological variation across donors while ensuring representation across all donors. Major cell classes from both datasets clustered together and cells from both datasets were uniformly distributed (Figure 1B-1D and S1A-S1C), except for samples from PCW 3-4 as it was contributed only by Zeng *et al*.’s dataset (Figures 1E; Supplementary Table 1).

**Figure 1:**
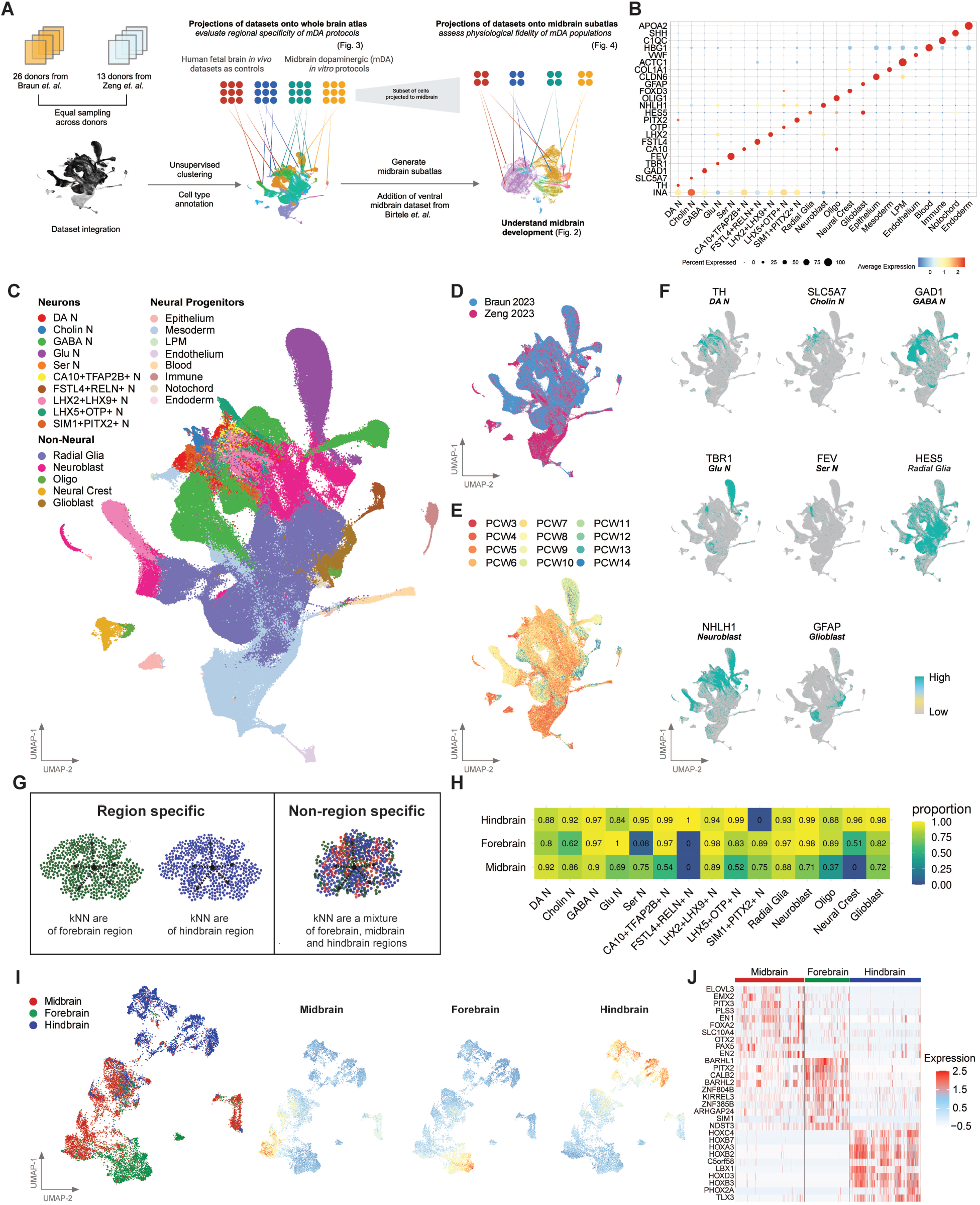
Construction of a fetal whole-brain single-cell atlas identified brain region-specific gene expression signatures across different neural cell types. (A) Schematic depicting the computational workflow for constructing the fetal whole-brain single-cell atlas and fetal midbrain subatlas, followed by their downstream applications. This includes projecting *in vitro*-derived datasets onto these reference atlases to evaluate their transcriptional fidelity. (B) Dotplot of gene expression of known marker genes across different annotated cell types in the fetal whole-brain atlas. Uniform Manifold Approximation and Projection (UMAP) plot of fetal whole-brain atlas showing (C) the annotated cell types, (D) the single-cell study, (E) the embryo age in post-conception week, and (F) the gene expression of known marker genes for selected cell types. (G) Schematic depicting the determination of region specificity by comparing the region identity of each cell’s k-nearest neighbors. (H) Heatmap showing the average proportion of k-nearest neighbors mapping to the corresponding brain region (midbrain, forebrain or hindbrain) across different cell types. UMAP plot of dopaminergic (DA) neurons showing (I) the brain region identity, (J) the midbrain region gene expression signature, (K) the forebrain region gene expression signature, and (L) the hindbrain region gene expression signature. (M) Heatmap of gene expression of top 10 marker genes, ordered by log2 fold change, for each brain region in DA neurons.

Next, we harmonized cell type labels across the reference datasets and curated a marker gene panel to annotate the integrated fetal whole brain atlas. Since our emphasis is on the development of neural cell types, we carried out a multi-step manual annotation of the atlas to distinguish neural and non-neural cell types before annotating finer neuronal subtypes. Our marker gene panel is derived from canonical markers supported by literature and cross-referencing to the original datasets’ annotation criteria (Figure 1B, 1F and S1D). This consensus annotation approach allowed us to define 23 cell types broadly classified as neurons, neural progenitors and non-neural (Figure 1B). To resolve neuronal subtypes, we extracted cells positive for pan-neuronal markers (*INA*+*DCX*+) and carried out subclustering at a higher resolution. We then applied a marker gene panel for the major subtypes of neurons such as *GAD1* for GABAergic neuron (GABA N) and *SLC5A7* for cholinergic neuron (Cholin N). The resulting annotation produced 10 subclasses of neurons, 5 groups which are classically defined by neurotransmitters and 5 other groups which do not fall into any of the categories and were labeled with their differentially expressed markers. Markers associated with the 23 cell types in the fetal whole brain atlas were reported (Supplementary Table 2).

Different brain regions comprise distinct cell types and microenvironments, which manifest as unique transcriptomic signatures. Here we utilize our integrated fetal whole brain atlas to identify cell types with region specificity and report the region-specific gene expression signatures for these cell types (Figures 1G-1J). To identify cell types with brain region specificity, we build a k-nearest neighbor (kNN) classifier by training it on region labels provided in Braun *et al*.’s dataset (Figure 1G). We focused on the 15 neural cell types for our classifier. Cell types with a region classification accuracy score higher than 0.65 were defined to be highly-region specific (Figure 1H). Of note, we observed high region prediction accuracy for cell types like midbrain DA neurons (score of 0.92) and forebrain glutamatergic neurons (score of 1.0), which are known to be distinctive neuronal types of the midbrain and forebrain respectively. In contrast, forebrain serotonergic neurons were low-scoring (0.08), suggesting a lack of forebrain-distinctive gene signature for these neurons. Interestingly, neural progenitor cell types (radial glia, neuroblasts and glioblasts) in our atlas displayed region-specificity for all 3 brain regions (Figure S1E).

Given that neurons and astrocytes are reported to have region specific gene expression, morphology and signaling, it stands to reason that their precursors may also display region specificity. The top differentially expressed genes (DEGs) distinguishing forebrain radial glia include well-known genes such as *DLX1* and *FOXG1*, while *EN1* marks midbrain radial glia and the *HOX* family of transcription factors are detected in the hindbrain radial glia gene signature.

In addition to well-established genes, we also report novel markers associated with each region-specific cell type (Figure 1I, 1J and Supplementary Table 3). Midbrain DA (mDA) neurons have been extensively characterized due to their selective vulnerability in Parkinson’s disease. In our atlas, we reported 21 genes that specify mDA neuron identity, many of which are well-corroborated by literature such as *EN1*^*16,17*^ and *PITX3*^18^. Of note, some novel markers potentially contributing to mDA development include *ELOVL3* (Very long chain fatty acid elongase 3) and *MYRIP* (myosin VIIA and Rab interacting protein).

### Fetal midbrain subatlas reveal midbrain-specific neuronal cell type development

Next, we set out to construct a subatlas focusing on the midbrain region. Unlike cortical development, human midbrain development remains relatively less studied. We used the region labels provided by Braun *et al*. to obtain cells of the midbrain region. To fully utilize our integrated whole brain atlas, we applied our region-based kNN classifier to cells from Zeng *et al*.’s dataset to infer the region identities for 10 neural cell types with region specificity scores of more than 0.65 (Figure 1H and 2A). We further augmented our fetal midbrain atlas with a recently published human ventral midbrain dataset by Birtele *et al*.^19^, creating a final midbrain subatlas comprising 102,335 cells (Figure 2B-2C and S2A; Supplementary Table 4). As far as we know, this is the largest developing midbrain-centric resource to date, spanning weeks 3 to 14 of human development.

**Figure 2:**
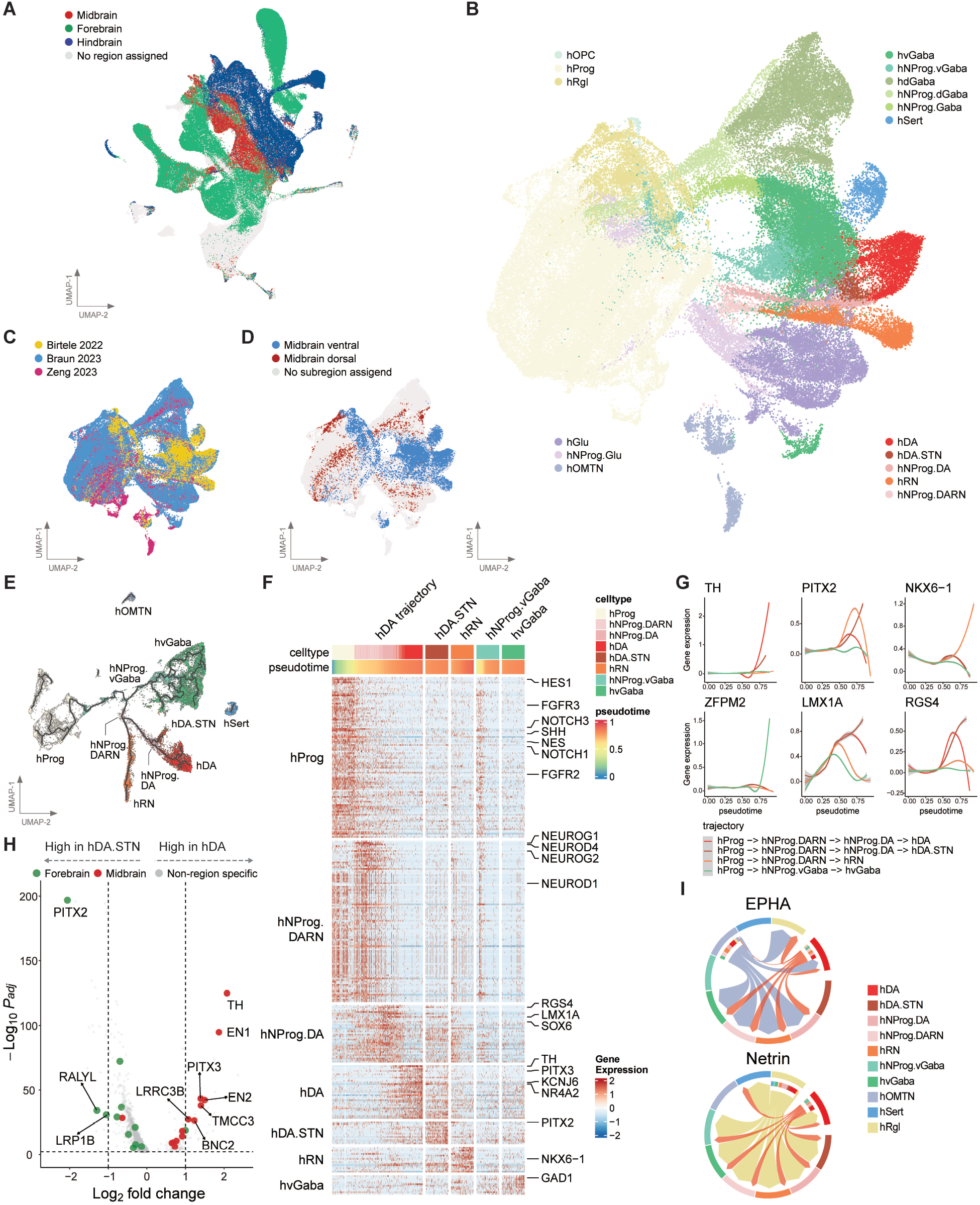
Fetal midbrain subatlas identified ventral/dorsal subregion-specific cell populations and a molecularly distinct subpopulation of hDA.STN neurons developmentally aligned to hDA neurons. (A) UMAP plot of fetal whole-brain atlas showing the brain region where cells are obtained. UMAP plot of fetal midbrain subatlas showing (B) the finer midbrain cell type annotations, (C) the single-cell study, and (D) the midbrain subregion i.e. ventral or dorsal. (E) UMAP plot of fetal midbrain ventral subregion-associated cells overlaid with Monocle3-inferred trajectories identifies four major trajectories giving rise to hDA, hDA.STN, hRN and hvGaba neurons. (F) Heatmap showing the gene expression of marker genes for the different cell types present in the four major trajectories identified by Monocle3 algorithm ordered by pseudotime. (G) Gene expression of selected genes across pseudotime that are upregulated in hDA (TH) or hDA.STN (PITX2) or hRN (NKX6-1) or hvGaba (ZFPM2) or hNProg.DA (LMX1A and RGS4). (H) Volcano plot showing the differential expression between hDA neurons and hDA.STN neurons. (I) Signaling patterns between fetal midbrain ventral subregion-associated cell types for the EPHA and Netrin signaling pathway.

To annotate the fetal midbrain subatlas, we adopted the cell type nomenclature defined by La Manno *et al*., 2016, in which they reported the first human ventral midbrain dataset and uncovered human specific molecular subtypes of various midbrain cell types. Our annotation gene panel consisted of marker genes calculated from the La Manno *et al*. dataset with some modifications (Figures S2B and S2C). Firstly, we added a “hGlu” cell type found in the dorsal midbrain compartment of Braun *et al*. dataset which was not present in La Manno *et al*’s ventral midbrain. Secondly, we collapsed certain cell types such as hRgl1, hRgl2a, hRgl2b, hRgl2c and hRgl3 into one common label “hRgl” (radial glia) if there was not enough difference between the subtypes in our midbrain atlas. Notably, we introduced novel midbrain cell type labels that were derived from a combination of our trajectory analysis and midbrain subregion information from the reference datasets. These include 6 subtypes of neuronal progenitors (hNProg [hNProg.Glu, hNProg.Gaba, hNProg.vGaba, hNProg.dGaba, hNProg.DARN and hNProg.DA]) which give rise to their corresponding neurons termed as glutamatergic neuron (hGlu), ventral GABA neuron (hvGaba), dorsal GABA neuron (hdGaba), red nucleus (hRN), dopaminergic neuron (hDA) and subthalamic nucleus neurons sharing developmental lineage with hDA (hDA.STN) (Figure 2B). The cell types hvGaba and hdGaba and their precursors hNProg.vGaba and hNProg.dGaba are labeled with a “v” or “d” subscript denoting their ventral or dorsal subregion bias based on midbrain subregion metadata (Figures 2D and S2E). Overall, the finer midbrain cell type annotation corresponds well with the major cell type categories of the whole brain atlas (Figure S2D).

To further delineate the developmental trajectories in the ventral subregion of the midbrain, trajectory inference was performed using the Monocle3 algorithm on ventral-associated cell populations in the fetal midbrain (Methods). The inferred trajectories suggest that hvGaba neurons diverges from hRN and hDA earlier in pseudotime (Figure 2E). Interestingly, hRN, hDA and hDA.STN neurons appear to share a common progenitor which we labeled as hNProg.DARN (Figure 2E). Anatomically, these three groups of neurons are in close proximity, supporting the idea that they may share a common intermediate progenitor during development. From our inferred lineages, hNProg.DARN then gives rise to hRN and hNProg.DA, the immediate precursor to hDA and hDA.STN (Figure 2E). Differential gene expression analysis on these ventral midbrain cell types identified well-known floor plate markers such as LMX1A and SOX6 expressed by hNProg.DA cells and classical dopaminergic marker genes TH and KCNJ6 for hDA (Figure 2F). We then looked into the temporal expression of selected markers for each ventral neuron subtype. Interestingly, LMX1A and RGS4 are highly expressed in both hDA and hDA.STN early in pseudotime, but hDA downregulates it later, while hDA.STN maintains high LMX1A expression (Figure 2G), supporting the idea that both neurons may share a common progenitor but diverge later by differential regulation of floor plate progenitor genes.

Detecting hDA.STN in our midbrain subatlas was surprising, given that the subthalamic nucleus is a diencephalic structure, located in the forebrain. Culturing dopaminergic neurons *in vitro* has been met with considerable cell heterogeneity despite high efficiency of initial patterning towards the floor plate identity. The subthalamic nucleus (STN) lineage has been reported to be closely related to the hDA lineage in mouse brain studies^20^. hDA.STN and hDA share classical mDA markers like *LMX1A* and *FOXA2*, making it challenging to distinguish in culture systems. We confirmed that *PITX2* is a robust marker of the STN lineage and reported more novel markers to aid with the distinction of these two closely related cell types (Figure 2H and Supplementary Table 5). Gene set overrepresentation analysis of differentially expressed genes revealed a predominance of axonogenesis and dopamine metabolism pathways in hDA neurons, while protein translation and protein folding related pathways were upregulated in hDA.STN neurons (Figure S2F). To understand intercellular communication differences between hDA and hDA.STN, we carried out cell-cell interaction analysis on ventral midbrain cell types. On a global level, hRN and hOMTN contribute to more outgoing interactions (Figure S2G). Both nuclei originate from the basal plate during development, a region that is highly neurogenic and may explain the strong signaling related to neuronal function such as Neuregulin (NRG) and Semaphorin pathways (Figure S2G and S2H)^21^. Comparing the signaling interactome of hDA and hDA.STN, we found the Ephrin-A (EPHA)^22^ and Netrin^23,24^ signaling pathways to be specific for hDA (Figure 2I). These pathways are essential for the formation of dopaminergic neuron circuits.

### Projection of midbrain differentiation protocols to integrated whole brain fetal atlas to assess fidelity of *in vitro* midbrain cultures

To determine the region and cell type of midbrain neuronal cultures, we retrieved published scRNA (single cell RNA) sequencing data related to midbrain patterning^25-36^. In total, we obtained 12 publicly available datasets and included our in-house midbrain organoid differentiation protocol to be evaluated. These include datasets that generate midbrain DA neurons both in 2D cultures and 3D organoid models. We broadly grouped them into two categories: 1) *in vitro* time series datasets and 2) Parkinson’s disease models. Here, we systematically evaluate each midbrain differentiation protocol for their brain region purity and DA neuron yield at a global scale by projecting first to our integrated whole brain atlas and at a finer resolution by projecting only cells with predicted midbrain identity to our midbrain subatlas (Figure 3A). To improve our cell type predictions, we implemented an additional filter by retaining only confident assignments of projected cell types determined by marker gene module scoring (Figure S3).

**Figure 3:**
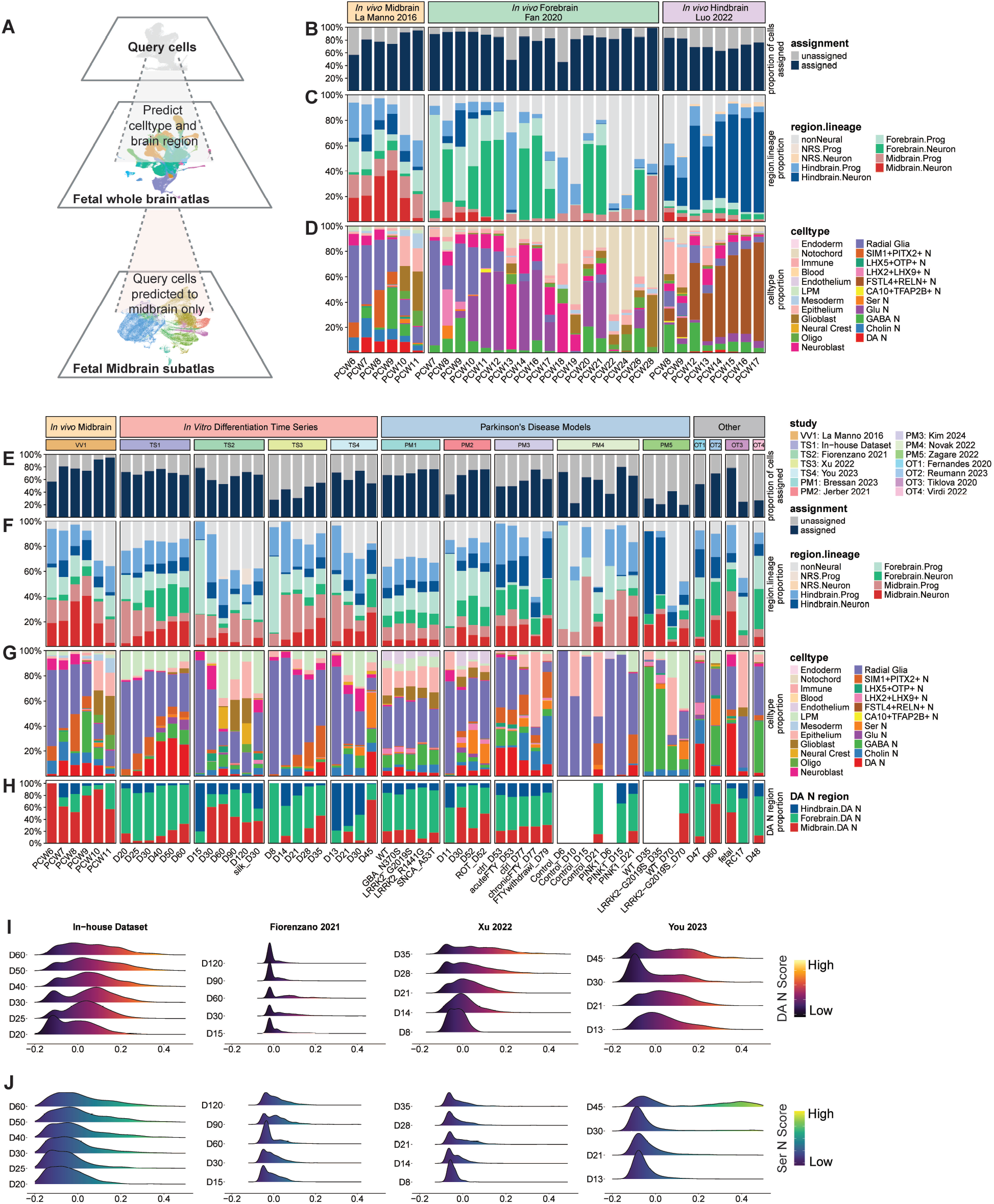
Projection of *in vitro*-derived midbrain datasets uncovered differences in cell type composition and brain region specificity across different *in vitro* differentiation protocols and Parkinson’s Disease models. (A) Schematic depicting the two-step mapping process of query datasets e.g. *in vitro* midbrain datasets onto our fetal whole-brain atlas and midbrain subatlas. Proportion plots of *in vivo* brain tissue single-cell studies across different developmental stages showing (B) proportion of cells confidently assigned, (C) region.lineage proportions for confidently assigned cells, and (D) cell type proportions for confidently assigned cells. Proportion plots of *in vitro*-derived single-cell datasets across different timepoints/conditions showing (E) proportion of cells confidently assigned, (F) region.lineage proportions for confidently assigned cells, (G) cell type proportions for confidently assigned cells, and (H) brain region proportions for confidently assigned DA neurons. Ridgeplot showing the distribution of (I) DA neuron gene expression signature and (J) Serotonergic neuron gene expression signature across different timepoints for selected *in vitro* midbrain differentiation time series studies.

We first demonstrate the reliability of whole brain projection using scRNA sequencing datasets of the developing midbrain^37^, cortex^38^ and cerebellum^39^ as controls for the midbrain, forebrain and hindbrain respectively. We performed the first-tier projection of these *in vivo* datasets to the integrated whole brain atlas and determined their proportion of assigned cell types and projected region identity. All *in vivo* datasets showed a high percentage of assigned cell types (Figure 3B) and the majority of their projected regions (>75%) belonged to their respective brain regions (Figure 3C). Among the projected cell types, we observed that the midbrain dataset has higher levels of DA neurons compared to datasets from other brain regions, while the forebrain dataset contains relatively more glutamatergic neurons (Figure 3D). The DA and glutamatergic neurons are typically found in the midbrain and forebrain respectively, confirming that the cell type predictions are as expected. Interestingly in the hindbrain dataset, we found high proportions of predicted FSTL4+RELN+ neurons. These may correspond to the cerebellar granule neurons which secrete reelin to guide Purkinje cell migration in the hindbrain^40^.

Next, we delve into the region projections of confidently assigned cells for each dataset (Figure 3E). We grouped the cell types into broad categories of progenitor, neuron and non-neural, and quantified the proportion projected to each brain region for each dataset (Figure 3F). Similar to the *in vivo* datasets, all *in vitro* queries consist of cells spanning across brain region identities. Strikingly, cells from forebrain and hindbrain regions comprise more than half of most midbrain datasets. We found that our in-house data and the dataset by Xu *et al*. displayed the highest yields of midbrain-predicted cells (Figure 3F). Among queries that have multiple timepoints, midbrain-predicted cells tend to peak between day 30 to 40 in culture^28,32,33^. Comparing 2D versus 3D culture conditions, it is not apparent which method is superior in terms of maximizing midbrain region purity. The 3D organoid protocols (in-house data and Fiorenzano *et al*., 2021) range between 25% to 35% midbrain regional identity at their respective peaks. 2D culture protocols can surpass this, such as the latest timepoint in Xu *et al*.’s dataset, although closer inspection reveals that most of the midbrain cells comprise of progenitors rather than mature neurons (Figure 3F). We also surveyed midbrain differentiation datasets generated for Parkinson’s disease modeling^27,29,30,34^. Some PD-associated mutants such as the LRRK2-G2019S mutant generated by Zagare *et al*. and the PINK1 mutant by Novak *et al*. show increased proportions of midbrain progenitors compared to their respective wild-type conditions. In contrast, other PD-associated mutants including GBA-N370S and SNCA-A53T did not show significant difference from the wild-type state, indicating that genetic mutations may not affect regional patterning in culture.

We next quantified the proportions of projected cell types for each dataset. Most protocols predominantly contain radial glia at early time points, and transition over time towards neuronal populations (Figures 3D and 3G). The neuronal distribution varied across protocols. Besides DA neurons, GABAergic and serotonergic neurons were observed in multiple datasets. Benchmarking against La Manno *et al*., it is clear that the DA neurons produced in culture are composed of a mixture of regions and their midbrain regional purity does not match that of dissected midbrain tissue (Figure 3H). Most midbrain protocols aim to generate DA neurons and minimize contaminants such as hindbrain serotonergic neurons. Hence, we scored the expression of mDA neuron and hindbrain serotonergic neuron gene modules in 4 query datasets that profiled their protocols over time. The peak of mDA gene module expression correlates with the peak in DA N yield in each protocol (Figure 3I). Likewise, we analyzed the hindbrain serotonergic neuron’s gene module scores of these time series protocols and found that 1 protocol contained substantially higher levels of hindbrain serotonergic gene signature compared to the rest, indicating that protocol may have excessive caudalization during *in vitro* patterning (Figure 3J).

### Fetal midbrain subatlas provides high resolution characterization of midbrain differentiation protocols

The projection pipeline we established allows us to further extract cells of midbrain regional identity and perform a second projection to the fetal midbrain subatlas (Figure 3A). With this approach, we are able to map to finer neural cell types introduced in the midbrain subatlas, without confounding factors from cell types of other regions. Eight datasets that contained more than 1000 cells after the first projection were mapped onto the fetal midbrain subatlas to obtain finer midbrain cell type annotations (Figure 4A). We then evaluated each protocol’s capacity to generate the different neuronal progenitors (hNProg) and neurons of the midbrain. We found that our in-house dataset generated the highest proportion of hNProg.DA within neuronal progenitors and hDA within neurons across the compared midbrain cultures. In contrast, most other datasets contained a substantial amount of hNProg.vGABA (>50%) and hNProg.Glu progenitors (>30%) (Figure 4B). Correspondingly, hvGABA and hGlu neurons were frequently observed in culture protocols (Figure 4C). We further calculated a ventral score for each dataset. Datasets that contained more ventral cell types such as hDA and hNProg.DA (Figure S2E) accordingly received higher ventral scores (Figure 4D).

**Figure 4:**
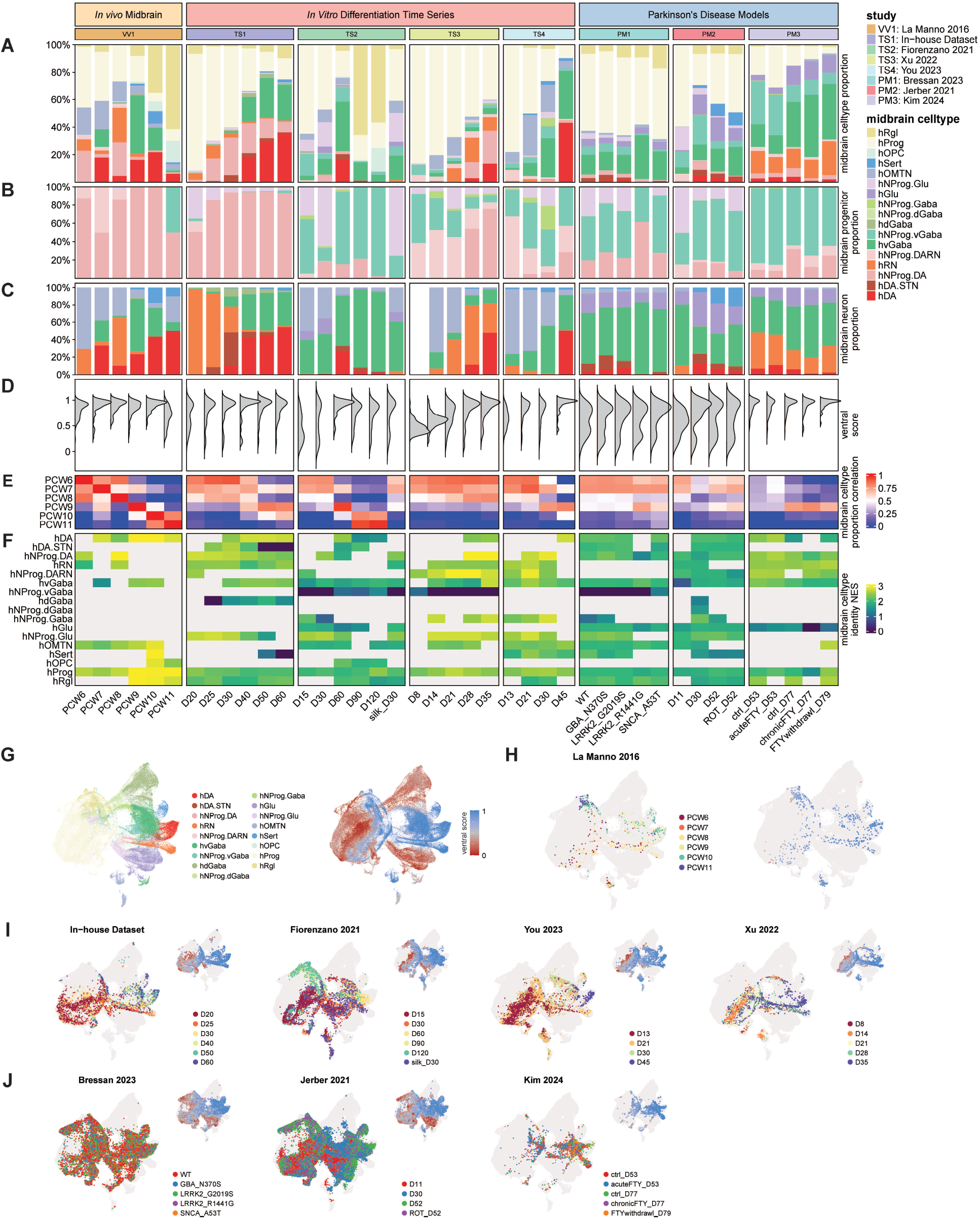
Secondary projection of midbrain-associated neural cells from *in vitro-*derived datasets revealed differences in cell type composition, ventral score of cells, and cell type identity strength across different *in vitro* differentiation protocols and Parkinson’s Disease models. Plots of *in vivo* and *in vitro* single-cell studies showing (A) midbrain cell type proportions, (B) progenitor proportions, (C) neuron proportions, (D) distribution of ventral score, (E) correlation of midbrain cell type proportions against *in vivo* timepoints, and (F) normalized enrichment score of cell type identity marker genes. (G) UMAP plot of fetal midbrain subatlas showing the midbrain cell type annotations and ventral score. (H) UMAP plot of *in vivo* midbrain data from La Manno *et al*. 2016 projected onto the fetal midbrain subatlas showing *in vivo* timepoints in post-conception weeks and ventral score. UMAP plot showing (I) different *in vitro* time series differentiation protocols and (J) Parkinson’s Disease models datasets projected onto the fetal midbrain subatlas showing their corresponding timepoints for *in vitro* differentiation time series and conditions for Parkinson’s Disease models. For panels I and J, the inset UMAP shows the ventral score. For panels H-J, cells from the fetal midbrain subatlas are plotted in a light gray color to provide context.

To assess how well each protocol recapitulates the cell type composition of the developing midbrain, we correlated their cell type proportions to each gestation week of the *in vivo* midbrain dataset (Figure 4E). Several datasets displayed high concordance with the developing midbrain in PCW 6 to 8, suggesting that *in vitro* cultures can recapitulate the cell type diversity in early development. However, most datasets did not show high correlation past PCW 8 despite continuing to mature *in vitro* up to 2 months, reflecting the challenges of generating neural cultures that resemble more mature states. Interestingly, one dataset that profiled midbrain organoids at day 120 showed high correlation to PCW 10 and 11, suggesting that it is possible to overcome this limitation given sufficient culture durations. To quantify the transcriptomic similarity between *in vitro* and primary midbrain cell types, we calculated cell type identity scores. We observed higher cell type identity scores for progenitor cell types (hProg and hNProg.DA) in the early time points of time series datasets, and higher cell type identity scores for neurons (hDA and hvGaba) at later experimental time points, in line with the appearance of these cell types in culture (Figure 4F). The distribution of projected cell types on the fetal midbrain subatlas’s UMAP space and ventral score distribution for each dataset is shown in Figures 4G-4J. Among time series datasets, we observe a progression from progenitor to neuron as expected (Figures 4I).

Overall, the two-tier projection method established in this study allowed for on-target region projection to the midbrain reference, an important step which facilitates the detection of rarer neural subpopulations in the midbrain. We comprehensively evaluated each midbrain differentiation protocol using a benchmarking panel of features unique to modeling midbrain development, such as their ventral score and similarity index to a developing fetal midbrain dataset. Our data shows that currently available midbrain differentiation protocols are varied in their approaches and outcomes to model the human midbrain. To enable future benchmarking of midbrain differentiation protocols, we present the fetal whole brain and midbrain atlases generated in this study as an online resource (BrainSTEM: http://brainstem.ouyanglab.com/) for users to explore and assess the atlases.

## Discussion

In this study we present the first integration of two large-scale developing human whole brain datasets. This integrated brain atlas allowed for the systematic identification of region-specific gene signatures, and demonstrated the variation in brain region specificity within key neural cell types. Including the dataset by Zeng *et al*. enabled us to profile the first trimester of brain development more completely as it uniquely contained samples from week 3 and 4 of gestation. Harmonizing both datasets during cell type annotation resulted in an integrated atlas with a better coverage of neuronal subtypes than each original atlas dataset alone.

A central contribution of this work involves the generation of the largest human fetal midbrain resource, created by extracting the midbrain cell types from the whole brain atlas and the addition of a ventral midbrain dataset. Within this curated midbrain subatlas, we further refined the cell type annotation, following the nomenclature on cell type labels introduced by La Manno *et al*. in 2016. The resulting fetal midbrain atlas consisted of novel neuronal progenitor cell types we identified through trajectory analysis and cell population proximity on UMAP space. Of note, we identified a small subpopulation of hDA (termed hDA.STN) bearing transcriptomic similarity to neurons of the subthalamic nucleus which are known to share common markers with dopaminergic neuron lineage. We believe that this cell population may have arisen from tissue dissection inaccuracies due to the anatomical proximity between the subthalamic nucleus and midbrain. Detecting hDA.STN in the midbrain subatlas suggests that our atlas is able to resolve rare cell types, given that it constitutes less than 1% in terms of cell numbers of the midbrain subatlas. Focusing on the ventral midbrain cell types, we identified genes that may distinguish key lineages of hDA and hRN neurons which originate from the floor plate and basal plate respectively (Fig. 2F). These findings offer new insights into the regulatory networks required for patterning within the midbrain, with implications for improving midbrain differentiation protocols that are enriched for hDA populations to be used for PD cell therapy.

Using the atlases we curated, we performed a comprehensive evaluation of available midbrain differentiation protocols, broadly categorized into either time series datasets or Parkinson’s disease models. Our approach to mapping *in vitro* brain data utilizes a biologically-driven framework (BrainSTEM) to first project datasets onto the whole brain reference then extract the on-target region (i.e. midbrain) to be mapped to the midbrain reference. This is in contrast to the conventional method of projecting midbrain datasets directly to a midbrain reference, which will prematurely limit the scope of the cell populations that can be mapped. We showed that, surprisingly, mapping forebrain or hindbrain datasets to a midbrain reference can still assign cell types relatively well (Figure S4A and S4B). This suggests that the computational mapping may have treated region-specific biological differences as technical differences that are “corrected” to arrive at a relatively good cell type assignment. These results clearly demonstrated the need for a biologically appropriate ground truth reference and relevant resolution to interrogate cell identity accurately. As neural culture systems are becoming more advanced, mapping these biologically complex *in vitro* products should reflect the diversity of cell type, regional and even sub-regional identities it may contain. With BrainSTEM, we can exclude more off-target regions as compared to directly mapping to a midbrain reference (Figure S4C-S4G). Importantly, the two-tier mapping of our BrainSTEM framework allows for rare cell populations such as hDA.STN to be mapped in the second tier of mapping which has increased resolution.

As neural culture systems become more complex and involve mimicking cells from different brain regions (e.g. assembloids), there is an increasing need to adapt computational practices to match the scale of the biological systems being interrogated. Conceptually, the multi-resolution projection method we propose is similar to iterative clustering, commonly used for cell type annotation in large datasets. Due to the hierarchical nature of cell type specification, cells may first be clustered at a coarse level into broad groups, before being sub-clustered into subtypes. As biological systems profiled at the single-cell level become more extensive, multi-level cell type annotation is becoming increasingly important. Similarly, a multi-resolution approach should be applied when projecting query datasets, such as midbrain culture single-cell data, onto reference atlases. Otherwise, mapping the query datasets directly onto a whole-brain atlas may lack the resolution needed to categorize finer cell types within each brain region. Conversely, direct mapping onto a midbrain reference may artificially inflate the number of confidently assigned cells, as non-midbrain-associated cells could be mapped spuriously, as demonstrated in our study. More importantly, the tiering needs to be biologically motivated. In our study, we only considered midbrain-associated neural cells in the second-tier mapping onto the midbrain subatlas, as non-neural cells (e.g., immune, mesoderm, and endoderm cells) do not display region-specific gene expression patterns. Thus, we envision that the BrainSTEM framework will pioneer the importance of multi-tier mapping to account for different cellular resolutions in complex biological systems.

Our comprehensive survey of currently available midbrain neuronal cultures revealed a wide range of midbrain cell types and proportions that constitute each dataset. While we observed a general trend towards maturation over time, temporal development of specific cell types appear different among protocols. In terms of cell type presence, PD cultures seem not to be different from healthy controls, although subtle distinctions may exist. Our analysis also highlighted the presence of numerous ‘off-target’ cells in all *in vitro* datasets, which may be erroneously identified as on-target cells thus rendering an inflated proportion if mapped directly. Lastly, as each dataset employs different culture protocols, our work provides a starting point for inferring morphogen effects on midbrain differentiation to enrich specific cell types like DA neurons, useful for improving PD cell therapy products.

### Limitations of the study

One important limitation of this study is the developmental time window that we have access to. The reference atlases span only the first trimester of development, and time points may have unequal coverage due to limited access to human fetal samples. In early works on the human and non-human primate embryo samples, DA neurons are mostly generated in the first trimester (between PCW 5-10) and start to send projections to targets in the striatum. Our atlases which cover this gestation window are well-positioned to profile the early stages of DA neuron development. However, DA neurons continue to undergo functional maturation and subtype specification well beyond the first trimester. Late maturation markers of DA neurons such as ALDH1A1 are accordingly not highly expressed in our atlases, limiting our ability to assess the maturation profile of *in vitro* midbrain protocols.

Increasing evidence supports the unique gene expression identity underlying regional differences among neural cell types. We have also identified putative marker genes acting in the midbrain for subregion specification. These novel marker genes are subject to experimental validation to confirm their spatial distribution and to determine the temporal expression profile during development. Through our survey of available midbrain differentiation protocols, we observed that each protocol can recapitulate some but not all aspects of the *in vivo* midbrain. Our fidelity metrics are certainly not exhaustive, future studies may consider profiling the epigenetic and spatial transcriptomic landscape of *in vitro* protocols given the emergence of multi-modal single-cell technologies. Nonetheless, the fetal midbrain subatlas represents a unified single-cell resource of the midbrain which can contribute to the conventional understanding of midbrain development *in vivo* and aid in refining midbrain protocols to advance the study of PD pathogenesis and cell therapy.

## Methods

### Metadata curation and preprocessing of human fetal whole brain scRNA-seq datasets

We used two recently published fetal brain datasets, Zeng *et al*. and Braun *et al*., to generate our fetal human whole brain atlas. These two reference datasets together span preconception week (PCW) 3-14 of gestation contributed by 39 donors. We sampled an equal number of cells (n = 17,665 cells) from each donor, except for two donors which had less cells than the rest, hence all cells were used from those two donors. All preprocessing and analysis was carried in Seurat (v4.9.9) unless indicated otherwise. Each down-sampled dataset was log-normalized using the ‘NormalizeData’ function, highly variable features extracted using ‘FindVariableFeatures’ function and scaled using ‘ScaleData’ function. Variance rank in each dataset was combined and we determined the top 3000 highly variable features from the combined rank. The consensus highly variable features were used for downstream analysis. We constructed the human fetal whole brain atlas by integrating the datasets using the Seurat anchor-based reciprocal PCA (RPCA) method^41^. Briefly, batch effect correction was performed using the 13 donors identifiers from Zeng *et al*. and on the 10x Chromium chemistry version (v2 or v3) from Braun *et al*. as the batches. Each batch was log-normalized, scaled and PCA was calculated on the top 3000 consensus highly variable features. Integration anchors were identified with the Seurat ‘FindIntegrationAnchors’ function and the top 50 PCs were used to integrate the datasets using Seurat ‘IntegrateData’ function specifying the Seurat RPCA method.

### Cell type annotation of the human fetal whole brain

Unsupervised clustering was performed using Seurat’s ‘FindNeighbors’ and ‘FindClusters’ functions using the top 50 PCs and a clustering resolution of 2.0. To annotate the clusters, we first harmonized cell type labels from the two reference datasets that compose our human fetal whole brain atlas. Matching cell types present in both datasets (e.g. “blood” and “erythrocyte”) were collapsed into the same label. Cell types present in only one of the datasets were retained (e.g. “Oligo” in Braun *et al*.). For neurons, we kept the subtype labeling by Zeng *et al*. (“Cholin N”, “DA N”, “GABA N”, “Glu N”) and added one more category for serotonergic neurons. A panel of marker genes informed by literature and databases for each cell type was used to annotate the atlas. To refine the clustering, we performed sub-clustering for selected cell types and benchmarked the annotation against our marker gene panel.

### k-nearest neighbor (kNN) classifier to identify cell types with distinct transcriptomic profiles across brain regions and assign region identity for these cell types

Brain region metadata is available in Braun *et al*. dataset, but not in Zeng *et al*. dataset. To determine the brain region specificity for each cell type, we trained a k-nearest neighbor (kNN) classifier with k = 10 on the top 50 PCs for each cell type to predict the brain region identity (midbrain / forebrain / hindbrain) at the single cell level, using the R package class (v7.3). This kNN classifier was trained on cells from Braun *et al*., leveraging on the available brain region metadata. Region specific cell types were defined as those with an average classification probability higher than 0.65 across all cells in that cell type. This indicates that the majority of each cell’s k-nearest neighbors belonged to the same brain region, implying distinct transcriptomic differences between brain regions for the particular cell type. As expected, all non-neural cell types have low classification probability. These cell types, together with neural cell types with classification probability lower than 0.65, were labeled as non-region specific (NRS) cell types. For the remaining region-specific neural cell types, the kNN classifier was then applied to cells from Zeng *et al*. to infer the brain region identity.

### Generation of the human fetal midbrain subatlas

The fetal midbrain subatlas comprises three reference datasets. Firstly, from the Braun *et al*. reference dataset, we subset all cells from the midbrain region using the region metadata provided. Secondly, from the Zeng *et al*. dataset, we used the kNN classifier to infer region identity for the 10 cell types that show midbrain region specificity, namely Cholin N, DA N, GABA N, Glu N, Ser N, LHX2+LHX9+ N, SIM1+PITX2+ N, Radial Glia, Neuroblast and Glioblast. From these cells, we kept those with inferred midbrain region identity. Lastly, we included a third reference human ventral midbrain dataset published by Birtele *et al*.. Each dataset was log-normalized and scaled separately before integration by Seurat RPCA approach. Top 3000 consensus highly variable genes were calculated by ‘SelectIntegrationFeatures’. All other parameters were the same as those in the fetal whole brain atlas construction.

### Cell type annotation of the human fetal midbrain subatlas

To annotate the integrated fetal midbrain subatlas, we first performed unsupervised clustering using Seurat’s ‘FindNeighbors’ and ‘FindClusters’ functions applied to the top 50 PCs and a clustering resolution of 3.0. We derived the midbrain marker gene annotation panel from La Manno *et al*.’s dataset and computed module scores for the top 10 markers of each cell type. Clusters were manually annotated based on the strength of module score expression. Acknowledging the transcriptional differences among neuronal progenitors (hNProg), we subset the hNProg cells with each of the terminal neuron population and generated UMAP to visualize the relationship between the hNProg and corresponding neuron cells. We then attached finer labels of hNProg subtypes to selected clusters based on their proximity to the corresponding neurons.

### Identification of differentially expressed genes

Differentially expressed genes were computed using Seurat’s ‘FindAllMarkers’ function in several analysis namely (i) identification of cell type specific genes for all the cell types in the fetal whole brain atlas, (ii) identification of brain region specific genes across the 10 cell types that show midbrain region specificity, (iii) identification of cell type specific genes for all the cell types in the fetal midbrain subatlas, (iv) identification of cell type specific genes across selected cell types in La Manno *et al*. dataset, (v) identification of cell type specific genes for all the cell types in the ventral subregion subset of the fetal midbrain subatlas and using Seurat’s ‘FindMarkers’ in (vi) identification of cell type specific genes between hDA and hDA.STN. Genes with average log fold change > 1.5, FDR < 0.01 were defined as highly expressed for that cluster.

### Reconstruction of developmental trajectories within fetal midbrain subatlas

As we are mainly interested in the developmental trajectories in the ventral subregion within the fetal midbrain, we further subsetted clusters with median ventral score of >0.5 across all the fetal midbrain cell types. This excluded dorsal-specific cell types e.g. hGlu and hdGaba. For the resulting ventral subregion subset, the top 50PCs were recalculated and then used to generate UMAP coordinates. To reconstruct the developmental trajectories, Monocle3^42^ (v1.3.7) was applied to the UMAP coordinates, identifying four major branches in the main trajectory, ending with the hDA, hDA.STN, hRN and hvGaba cell type respectively. To calculate the pseudotime, the diffusion pseudotime from scanpy package (v1.10.1) was used, setting the single cell in cluster 36 (which has the lowest median ventral score) that is furthest away from all other clusters as the root cell i.e. the cell with zero pseudotime. The top marker genes identified using Seurat’s ‘FindAllMarkers’ function are then visualized across the four different branches along pseudotime. Furthermore, the gene expression pattern of selected genes across the four different branches along pseudotime are interpolated using the loess function in R.

### Cell-cell interaction analysis

Cell-cell communication signaling pathways in the midbrain atlas were analyzed against the curated repositories of ligand-receptor interactions from CellChatDB^43^ (v1, v2) and CellPhoneDB^44^ (v4.1). Communication probabilities accounting for sample variability using the default pipeline and interactions found in at least one study were kept. ‘netAnalysis_signalingRole_heatmap’ was used to visualize the relative signaling strengths for all significant signaling pathways. ‘netVisual_chord_cell’ was used to visualize the relative signaling strengths between interacting cell types of selected pathways that are specific to hDA or hDA.STN cell type.

### Projection of query datasets to the fetal brain reference atlases

To compare midbrain differentiation protocols to the fetal whole brain reference, we used Seurat’s ‘FindTransferAnchors’ with RPCA approach, followed by ‘TransferData’ functions with default settings. For each query, we calculated a module score for each predicted cell type using the whole brain reference’s top 20 marker genes. Cells with a module score > 0 for their predicted cell type were retained as confidently assigned cells. Next, to assign region identity, we plotted the module score of each regional cell type’s top 20 marker genes. The region with the highest module score is taken to be the predicted region of the query cell.

After the first projection to the fetal whole brain atlas, we subset the midbrain region of each query dataset. The midbrain regions from selected datasets were used for the second mapping to the fetal midbrain subatlas. Additionally, datasets containing fewer than 1000 cells retained were discarded. We utilized Seurat’s ‘MapQuery’ function, a wrapper function of ‘TransferData’, ‘IntegrateEmbeddings’ and ‘ProjectUMAP’ to get predicted cell type identities of each query and their projected coordinates on the reference UMAP.

### Comparison of query datasets

To compare query datasets after the first projection to the whole brain, we computed the (i) proportion of cells confidently assigned, (ii) region.lineage proportions for confidently assigned cells, (iii) cell type proportions for confidently assigned cells, and (iv) brain region proportions for confidently assigned DA neuron cells. To compare query datasets after the second projection to the midbrain subatlas, we computed the (i) midbrain cell type proportions, (ii) progenitor proportions, (iii) neuron proportions, (iv) distribution of ventral score, (v) correlation of midbrain cell type proportions against *in vivo* timepoints, and (vi) normalized enrichment score of cell type identity marker genes. For ventral score calculation, see the next section. To correlate the midbrain cell type proportions, we first calculated the midbrain cell type proportions for *in vivo* midbrain La Manno *et al*. dataset for each timepoint spanning PCW 6-11. The Pearson correlation was then calculated between each of these *in vivo* timepoints and the cell type proportions for confidently assigned cells in each timepoint/condition for the query datasets. To assess the transcriptional robustness of the assigned cells, we first calculated the log2-fold-change for all genes comparing the predicted cell type of interest against all other predicted cell types in the query dataset. This log2-fold-change was then used as an input for Gene Set Enrichment Analysis (GSEA) using the top 50 marker genes for the predicted cell type of interest as the gene set. The resulting Normalised Enrichment Score reflects how well the upregulated genes in the predicted cell type of interest are enriched for the marker gene set.

### Ventral score classification

The R package ranger (v0.15.1) was used to train a random forest classifier on midbrain ventral and dorsal subregion differences based on labels provided by Braun *et al*. dataset and Birtele *et al*. dataset. The model was used to predict a score for each query at a single cell level, with higher scores corresponding to higher ventral identity. The resulting ventral score was visualized on UMAP using Seurat’s ‘FeaturePlot’ function.

## Resource Availability

## Lead Contact

Further information and requests for resources should be directed to and will be fulfilled by the lead contact, Alfred X. Sun

## Materials availability

This study did not generate new reagents.

## Data and code availability

Raw or pre-filtered UMI count matrices for the datasets used in this study were downloaded either from GEO repositories or obtained directly from authors (see Table 1 for more details). The processed Seurat object for the fetal whole-brain atlas and midbrain subatlas will be made available at the time of publication. The data can also be browsed interactively online at http://brainstem.ouyanglab.com/. An R package named ‘BrainSTEM’ to allow users to map their own midbrain culture single-cell datasets will be made available at the time of publication. All data were analyzed with commonly used open-source software programs and packages as detailed in the Methods section. The analysis code will be publicly available on GitHub at the time of publication.

## Acknowledgements

This work is supported by National Research Foundation Fellowship (NRF-NRFF14-2022-0008, to AXS), Singapore PARKinson’s disease Translational Clinical Programme (MOH-OFLCG18May-0002, to Tan Eng King and AXS), USyd-NUS Ignition Grant (2024USYD-NUSP01, to PY and AXS) and Singapore National Medical Research Council (NMRC) under OF-YIRG funding (MOH-OFYIRG21nov-0004, to JFO). Schematic diagrams (Figure 1A, 1G and 3A) were generated using Biorender (biorender.com).

## Author Contributions

HSYT and LX collected and retrieved scRNA-seq data of midbrain differentiation datasets used for mapping. LX performed data preprocessing and integration using the pipeline developed with the help of JFO for the whole brain and midbrain fetal atlases. LX, HSYT and JFO did cell type annotation for both atlases, with support from AXS. LX used the kNN classifier to determine cell types with region specificity and HSYT calculated region-specific markers for the region-specific neural cell types. JFO performed pseudotime analysis on the ventral-associated cell populations in the midbrain subatlas. CC and PY performed cell-cell interaction analysis. LX carried out the two-tier mapping on all curated single-cell query datasets and calculated cell type module scores to retain confidently assigned cells with support from HSYT, JFO and AXS. HSYT calculated midbrain DA neuron and hindbrain serotonergic neuron module scores for time series datasets. JFO developed the ventral score classifier and LX implemented it on all query datasets which were mapped to the midbrain subatlas. HSYT, LX, JFO and AXS conceived the project. HSYT and JFO wrote the manuscript with input from all coauthors. All authors read and approved the manuscript.

## Figure Captions

**Supplementary Figure 1: Additional plots related to the construction of the fetal whole-brain single-cell atlas**. (A) Proportion plot showing the proportion of cells coming from each of the two single-cell studies for each cell type. UMAP plot of fetal whole-brain atlas showing the original cell type annotation from (B) Braun *et al*. and (C) Zeng *et al*. (D) Dotplot of gene expression of full marker gene lists across different annotated cell types in the fetal whole-brain atlas, related to main Figure 1B. (E) UMAP plot of different region-specific neural cell types showing the brain region where the cells are collected as well as the midbrain / forebrain / hindbrain region gene expression signature, related to main Figure 1I and 1J.

**Supplementary Figure 2: Additional plots related to the construction of the fetal midbrain subatlas**. (A) Proportion plot showing the proportion of cells coming from each of the three single-cell studies for each midbrain cell type. Dotplot showing (B) the average module score across each midbrain major cell type and (C) gene expression of full marker gene lists across different midbrain cell types. (D) Alluvial plot showing the concordance in annotation from whole brain cell type to midbrain cell type. (E) Proportion plot showing the proportion of cells obtained from different midbrain subregions across different midbrain cell types. (F) Functional analysis of Gene Ontology pathways associated with differentially expressed genes between hDA and hDA.STN cell type. (G) Heatmap of outgoing and incoming signaling patterns from cell-cell communication analysis, across different midbrain cell types. (H) Selected signaling pathways showing different patterns between hDA and hDA.STN cell type.

**Supplementary Figure 3: Additional plots related to the projection of *in vitro*-derived midbrain datasets onto fetal whole-brain single-cell atlas**. Series of dotplots before module score filtering (left panel) and after module score filtering (right panel) across predicted cell types for each of the projected *in vivo* and *in vitro*-derived midbrain datasets.

**Supplementary Figure 4: Plots demonstrating the limitations of direct mapping and advantages of the two-tier mapping procedure introduced in this study**. Heatmap showing the proportion of cells for different cell type labels predicted using direct mapping, normalized across each cell type label predicted using two-tier mapping for (A) *in vivo* forebrain dataset Fan 2020 and (B) *in vivo* hindbrain dataset Luo 2022. (C) Venn diagrams showing the overlap of DA neurons predicted using two-tier mapping and direct mapping for *in vivo* midbrain and different *in vitro* differentiation protocols datasets. (D) Brain region (midbrain / forebrain / hindbrain) gene expression signatures calculated for DA neurons predicted using (i) two-tier mapping (TTM), direct mapping (DM) and cells uniquely called from direct mapping (DM only) for *in vivo* midbrain and different *in vitro* differentiation protocols datasets. Proportion plot showing (E) the predicted cell type label using different mapping approaches and (F) the predicted brain region (midbrain / forebrain / hindbrain) for *in vivo* midbrain and different *in vitro* differentiation protocols datasets. (G) Schematic venn diagram illustrating the definition of TTM, DM and DM only labels shown in Figure S4D-S4F.

