## Supplementary figures and images for "BrainSTEM: A multi-resolution fetal brain atlas to assess the fidelity of human midbrain cultures"

### Supplemental Figures

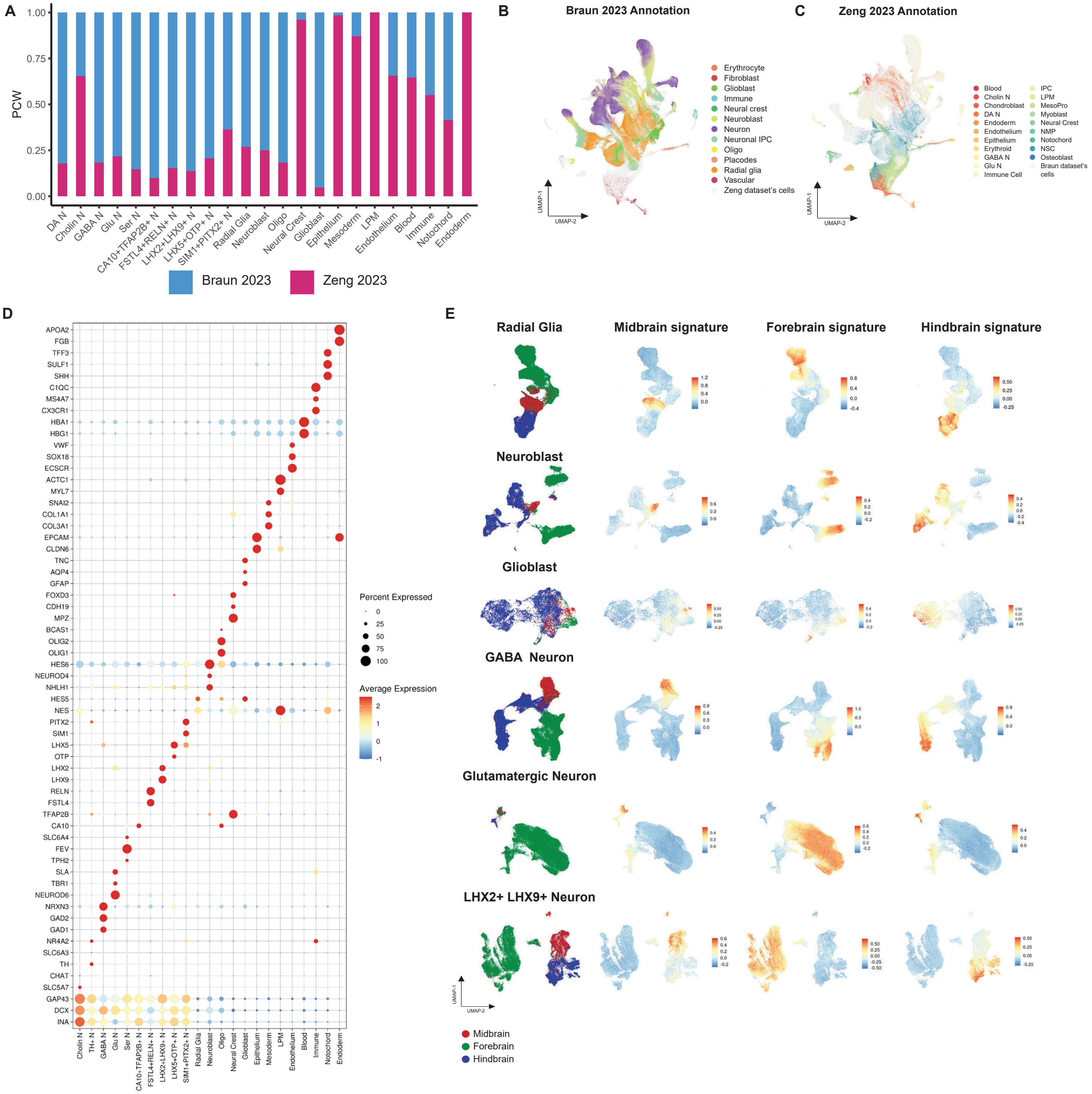

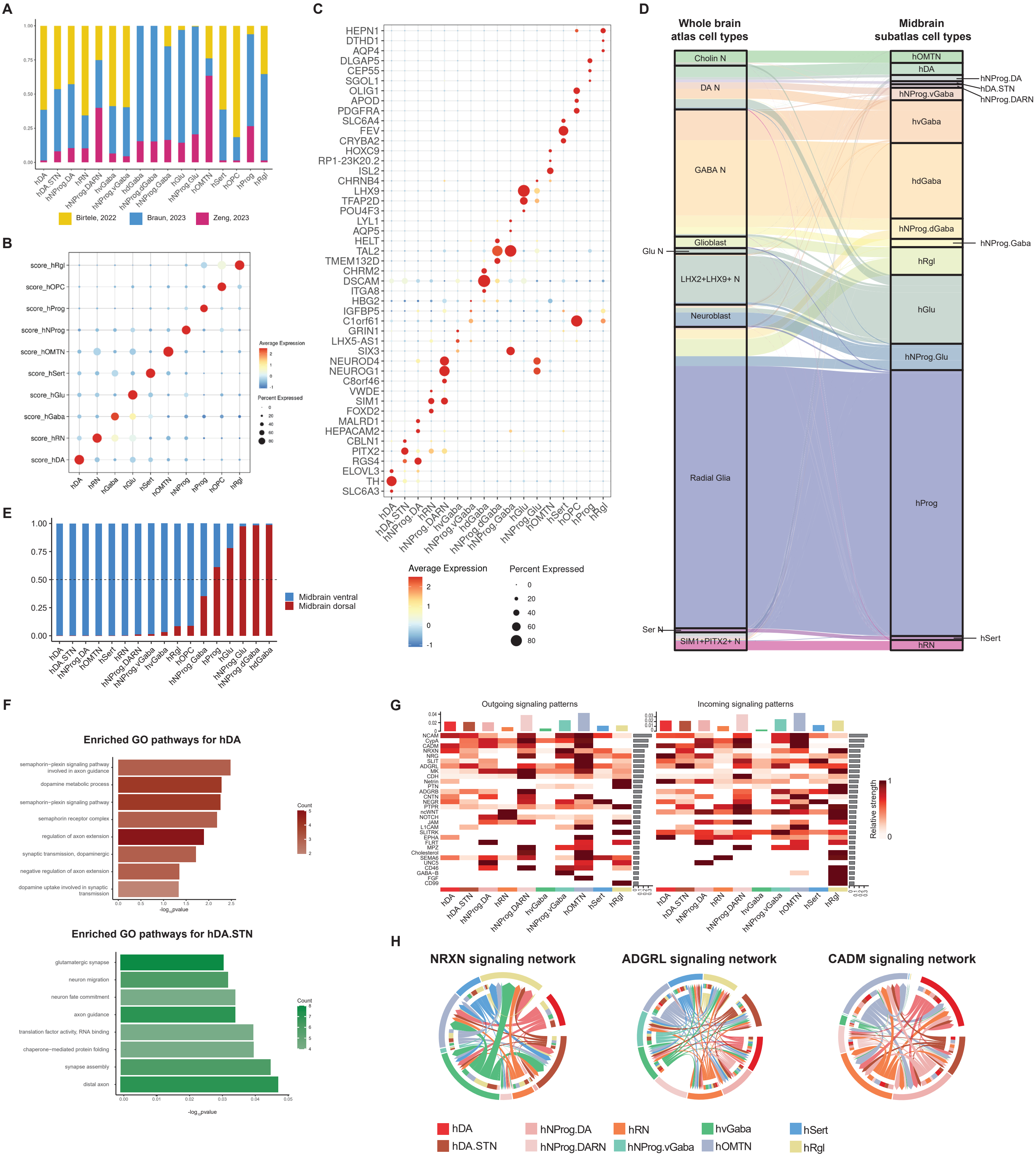

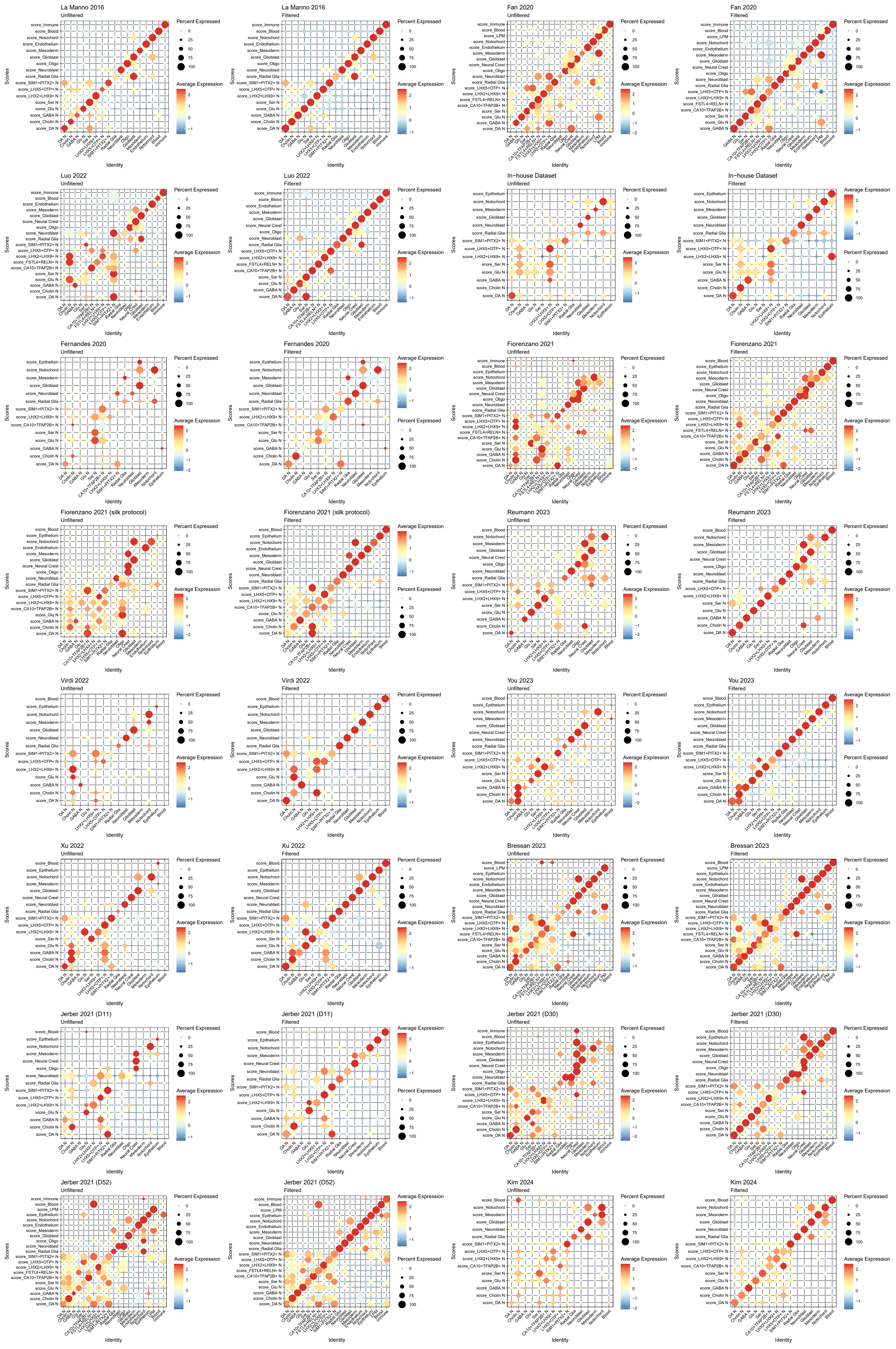

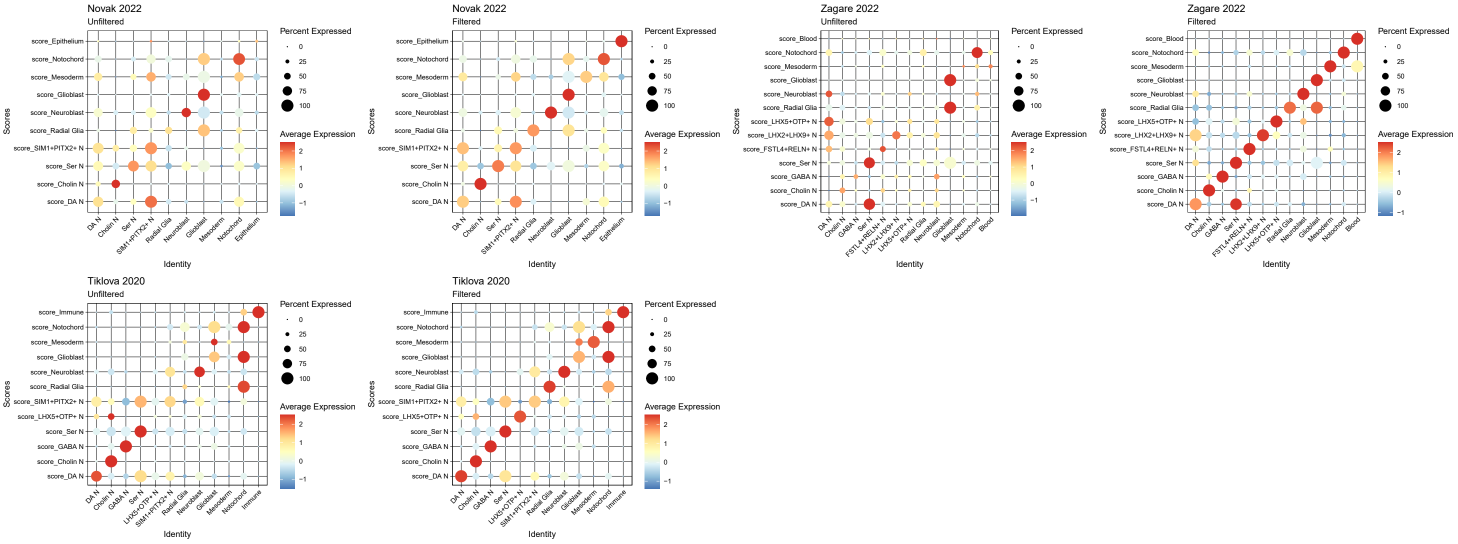

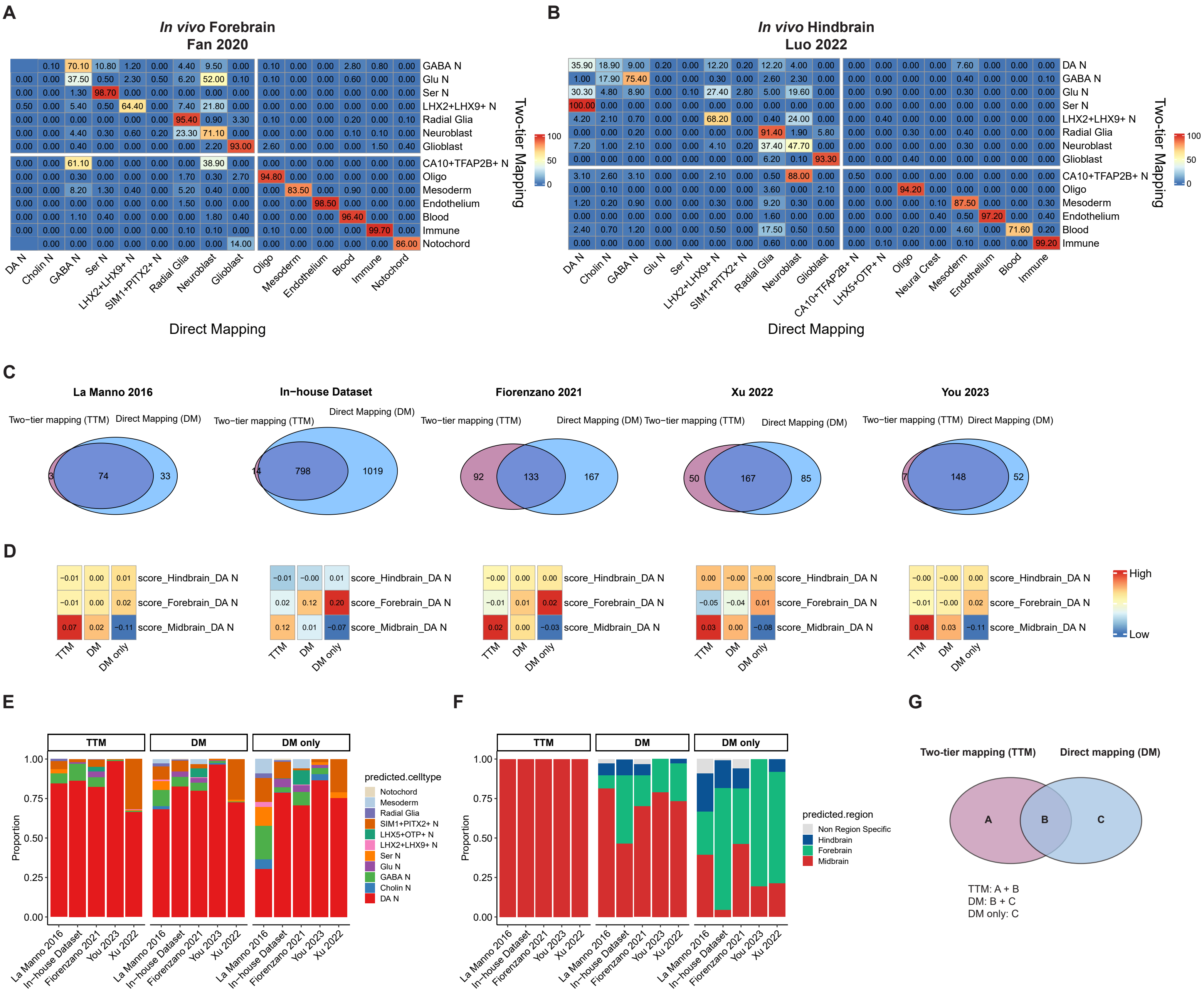
